# Soft multimodal opto-electric biointerfaces for co-localized optical and electrical recording of cell function

**DOI:** 10.1101/2022.12.21.521519

**Authors:** Sofian N. Obaid, Nathaniel Quirion, Jade Balansag, Nicolas Daza, Xinyu Shi, Zhiyuan Chen, Luyao Lu

## Abstract

Optical fluorescence and electrical monitoring of cell activity are two powerful approaches to study organ functions. Simultaneous recording of optical and electrical data types will provide complementary information from and take advantage of each approach. However, devices that can concurrently record optical signals from the same cell population underneath the microelectrodes have not been widely explored and remain a grand technical challenge. This work presents an innovative flexible opto-electric device that monolithically integrates transparent gold nanogrid microelectrodes directly above microscale light-emitting diodes, photodetectors, and optical filters to achieve co-localized crosstalk-free optical fluorescence and electrical recording. The optimized gold nanogrid microelectrodes show excellent optical transparency (>81%) and low normalized 1 kHz electrochemical impedance (6.3 Ω cm^2^). The optical recording subsystem offers high wavelength selectivity (>1,300) and linearity (R^2^ >0.99) for exciting and capturing green fluorescence from various fluorescent reporters in measurement ranges relevant to *in vivo* applications with minimal thermal effects. The opto-electric device exhibits remarkable durability under soaking for 40 days and repetitive mechanical bending for 5,000 cycles. The work may provide a versatile approach for constructing mechanically compliant biointerfaces containing crosstalk-free optical and electrical modalities with widespread application potentials in basic and clinical research.

## INTRODUCTION

Electrical recording is the gold-standard in monitoring action potentials, extracellular field potentials, and studying the functions of cells, tissues, and organs with high temporal resolution.^1–3^ Despite the tremendous impact of classic electrical recording in basic and applied biomedical research, it is unable to readout important functional parameters such as calcium dynamics or target specific types of cells. Optical recording is complementary and orthogonal to electrical recording^4–6^ A list of essential cellular parameters can be measured optically using appropriate fluorescent dyes, such as membrane potentials, calcium dynamics, metabolic states, and neurotransmitter release.^7–12^ The use of genetically encoded fluorescent reporters further allows optical recording from different cell types.^13–14^ Despite those advantages, the temporal resolution from optical recording is low due to the chemical properties and dynamics of currently available fluorescent reporters. As a result, simultaneous monitoring of optical fluorescence and electrical signals at the same device-tissue interface is highly desirable to leverage the advantages of both technologies.

Multimodal devices that offer the capabilities for simultaneous optical and electrical sensing are still in relatively early stages of development and are generally more difficult to implement than single function recording tools. A few devices that assemble opaque electrodes with optical fibers have been reported.^15–17^ However, these devices have certain deficiencies. For example, the electrodes in those devices are tens of micrometers away from the tip of the fibers since the opaque electrodes will block the direct light delivery to and collection from the areas beneath the electrode sites. In addition, these opaque electrodes will produce severe photoelectric artifacts that scale with optical power.^18–20^ Thus, those devices prevent co-localized optical and electrical recording from the same cells to fully understand how the two data types are related to each other. In addition, for future wireless implantable operations, on-chip microscale photodetectors (μ-PDs) and microscale light-emitting diodes (μ-LEDs) are a better choice than optical fibers, which can otherwise significantly restrict animal movements or lead to fiber entanglement during social behaviors.^21–22^ In this context, an ideal multimodal device will (1) contain internally mounted pairs of microscale optical and electrical sensing components that can operate under fully implanted condition; (2) match the location of the microelectrode and photodetector/light source for co-localized optical and electrical recording in a crosstalk-free manner; (3) exhibit excellent flexibility for conformal integration with soft tissues/organs and reduce the mechanical mismatch at the device and biological interface.

The rise of transparent microelectrodes has allowed efficient light delivery through the microelectrode areas for optical and electrical investigations at the same site with suppressed photoelectric artifacts.^23–27^ Advances in microfabrication technologies have enabled attachment of separately fabricated indium tin oxide (ITO)^28^ or graphene^29^ transparent microelectrodes and blue μ-LED probes for co-localized electrical recording and optogenetic modulation. Unfortunately, some critical shortcomings exist in these device fabrication. For instance, manual bonding different probes with each other using adhesive/glue to accommodate the additional modalities will increase the dimensions of the resulting devices, which results in more severe tissue damage for implantable applications. Misalignment for the microscale components on different probes can also occur in such a fabrication process when the sizes of the microelectrodes and μ-LEDs are reduced. Meanwhile, optogenetics only requires μ-LEDs as light sources to excite opsins while for fluorescence recording, μ-LEDs with additional μ-PDs and effective optical filters are needed to not only excite but also selectively record fluorescence from the fluorescent reporters. As a result, the development of opto-electric devices for on-chip, co-localized synchronous multimodal optical fluorescence and electrical sensing is still challenging. Further advancements on device schemes are highly desirable to realize the full potential of the multimodal recording technologies.

We recently reported single function probes integrating μ-PDs and μ-LEDs for *in vivo* calcium fluorescence recording,^21–22^ and strategies to integrate metal grid transparent microelectrodes precisely on the surface of μ-LEDs for co-localized optogenetic modulation and electrophysiology.^30–31^ In this work, we take a major leap forward by designing, fabricating, and characterizing a flexible and multimodal opto-electric probe that monolithically integrates (1) a μ-LED for on-chip excitation of the widely used green fluorescent reporters (GFRs); (2) a μ-PD coated with an effective optical filter for direct on-chip measurement of the resulting fluorescence signals from the GFRs; (3) a gold (Au) nanogrid microelectrode patterned ontop of the optical components for co-localized on-chip capturing of electrical signals. The nanogrid microelectrode is vertically stacked above the μ-LED, μ-PD, and optical filter where the fluorescence excitation and emission light can pass through for simultaneous optical fluorescence and electrical recording from identical cells. The detailed microfabrication process to build such a system is described, which combines photolithography, electron beam lithography (EBL), and micro-transfer printing techniques. The systematic characterizations of the electrochemical, mechanical, optoelectrical, and thermal properties validate the excellent performance of the device. Experimental results show that the Au nanogrid features a transmittance >81% from 400 to 800 nm and a low normalized 1 kHz electrochemical impedance of 6.3 Ω cm^2^ while the μ-LED and μ-PD/optical filter pair exhibits a high wavelength selectivity (>1,300) and an excellent linearity (R^2^ >0.99) for detecting weak green light signals. Temperature increase from the μ-LED during optical recording is carefully investigated for safe *in vivo* operations. Soaking test in a phosphate-buffered saline (PBS) for up to 40 days and cyclic mechanical bending test against a 5 mm radius for up to 5,000 cycles reveal its durability and flexibility for chronic applications. Finally, benchtop tests demonstrate the suitability of the opto-electric device to record programmed electrical waves and fluorescence signals from various GFRs. The results in this work shed light on the possible use of the multimodal flexible opto-electric biointerfaces with desirable characteristics for simultaneous on-chip optical fluorescence and electrical recording of the same cell populations, with broad potentials in many areas of basic and translational biomedical applications.

## MATERIALS AND METHODS

### Design and Fabrication of the Multimodal Opto-electric Device

Figure 1a shows a schematic illustration of the unique device architecture, fabrication process, and integration strategy for the multimodal opto-electric probe. The purpose is to monolithically integrate a μ-LED, a μ-PD, an optical filter, and a Au nanogrid transparent microelectrode on a flexible substrate for co-localized optical and electrical recording. The fabrication process began by spin coating polydimethylsiloxane (PDMS, Sylgard 184, Dow Corning, base:curing agent = 10:1) on a glass slide at 2,000 rpm for 50 s as the adhesive layer. The glass slide serves as the rigid handling substrate for microfabrication. A 25 μm thick transparent and biocompatible polyethylene terephthalate (PET) film (CS Hyde Company) was laminated on PDMS as the flexible substrate for the opto-electric device. A 7 μm thick SU-8 2007 (MicroChem Corp.) was spin coated on the PET film at 3,000 rpm for 40 s, exposed by ultraviolet (UV) light, and cured at 110 °C for 15 min as the planarization layer. A metallization process using photolithography (AZ® nLOF 2070 photoresist, Integrated Micro Materials), electron beam (e-beam) evaporation, and liftoff in acetone defined the Cr/Cu (20/100 nm) cathodes, anodes, and interconnects for the μ-PD (300 × 300 × 100 μm^3^, TCE12-589, Three Five Materials Inc.) and the μ-LED (220 × 270 × 50 μm^3^, C460TR2227-0216, Cree Inc.). The μ-LED emits blue light with the peak at 462 nm,^31^ which is suitable to excite the widely used genetically encoded calcium indicator GCaMPs (excitation/emission maxima at 488/512 nm),^32^ voltage indicator ASAP3 (excitation/emission maxima at 488/520 nm),^33^ neurotransmitter indicator GACh (excitation/emission maxima at 488/520 nm),^34^ and the endogenous fluorescent marker of cellular energy metabolism flavin adenine dinucleotide (excitation/emission maxima at 470/525 nm).^35^ The blue μ-LED and μ-PD were micro-transfer printed onto the Cr/Cu cathodes and anodes using soft lithography defined PDMS elastomeric stamps. The central recessed structures of the stamps match the sizes of the μ-LEDs and μ-PDs. A reflow soldering process with an In/Ag solder paste (Indalloy 4, Indium Corporation) at 150 °C for 3 min formed the robust mechanical and electrical connections. Our previous experimental and Monte Carlo simulation results show that the major fluorescence measurement volume for on-chip optical sensors is at the direct interface between the μ-PD and μ-LED.^21^ As a result, the μ-LED and μ-PD were placed next to each other to form a high performance optical fluorescence recording pair. Absorber dye ABS 473 (Exciton) was mixed into a transparent SU-8 2007 photoresist at different weight ratios, spin coated, and photolithographically patterned on the μ-PD as the optical filter for blue light. Another 150 μm thick SU-8 layer was spin coated, UV exposed, and fully cured ontop of the μ-LED and μ-PD at 110 °C for 10 min as the passivation and encapsulation layer for the optoelectronic components.

**Figure 1.**
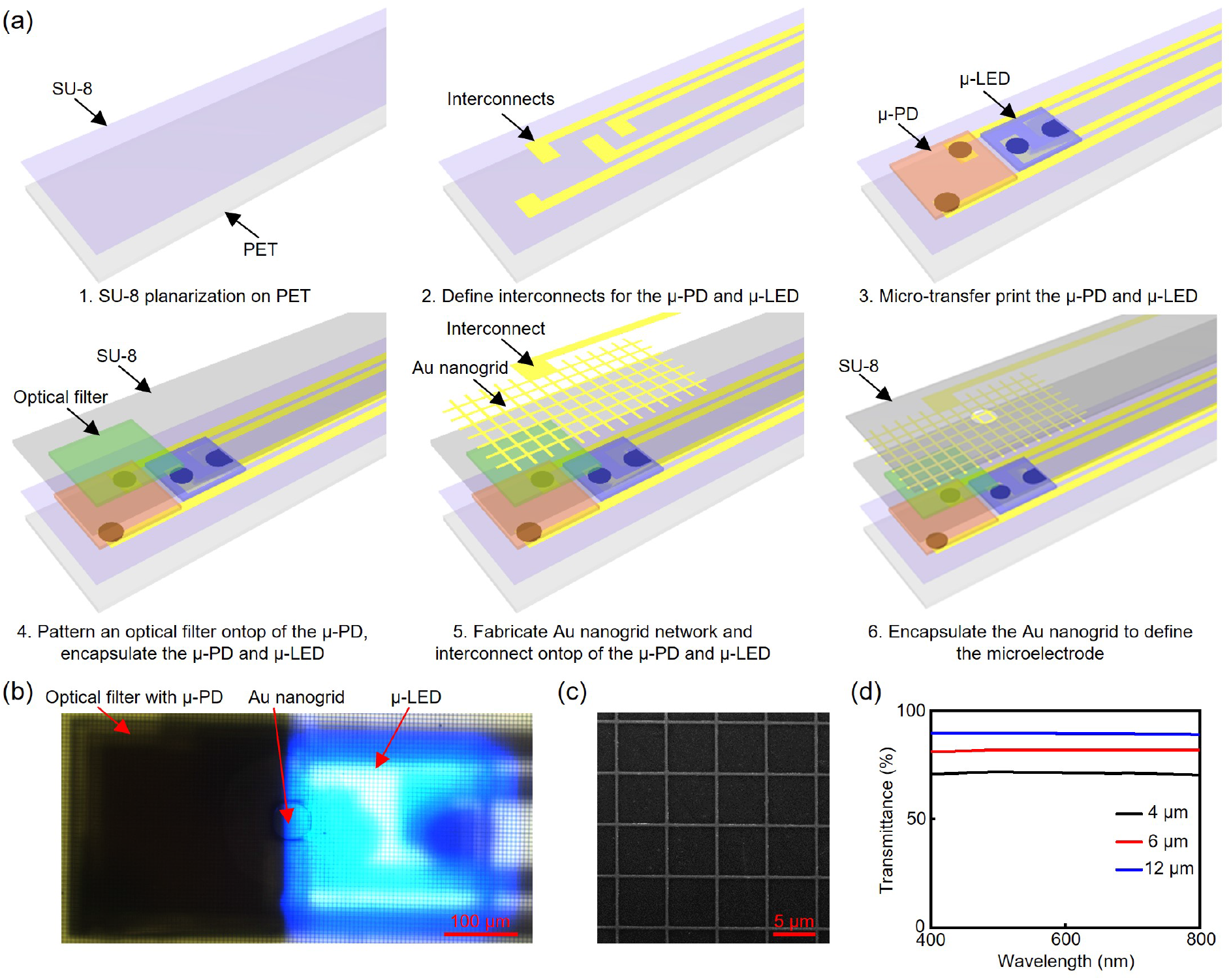
Schematic illustration of the flexible multimodal opto-electric device that combines microscale optoelectronics and transparent Au nanogrid microelectrodes. (a) Schematic fabrication scheme of the flexible multimodal opto-electric device. (b) Optical image of a flexible multimodal opto-electric device containing a blue μ-LED, a μ-PD with optical filter and a transparent Au nanogrid microelectrode. (c) SEM image of the Au nanogrid network in (b). (d) Transmittance spectra of Au nanogrid films with 400 nm nanogrid width, 40 nm thickness, and different pitch values at 4, 6, and 12 μm, respectively.

The device was then prepared for EBL with PMMA A4 resist (MicroChem Corp.). The Au nanogrid patterns were defined by 10 kV, 60 μm aperture beam with a current of 0.75 nA, and a dose of 150 μC/cm^2^. Fine alignment of the nanogrid patterns ontop of the optoelectronic components during EBL was achieved using the Cr/Cu metal interconnects for the μ-LEDs and μ-PDs as markers. A bilayer Cr/Au (20/40 nm) was deposited by e-beam evaporation. Liftoff in acetone completed the Au nanogrid microelectrode fabrication. A top SU-8 2007 layer defined the 50 μm diameter microelectrode window and encapsulated the rest of the device via photolithography. The microelectrode window overlapped with the μ-LED and μ-PD interface so the positions of the optical fluorescence and electrical recording sites are precisely matched. Laser cutting determined the geometry of the opto-electric probe. Delaminating the device from the glass slide handling substrate completed the fabrication and integration process. Figure 1b shows an optical image of a final multimodal probe with all optical and electrical components and the blue μ-LED on.

### Electrochemical Measurements

Electrochemical Impedance Spectroscopy (EIS) and cyclic voltammetry (CV) of the Au nanogrid microelectrodes were measured with a Gamry Reference 600+ potentiostat/galvanostat/ZRA (Gamry Instruments Inc.) employing a three-electrode configuration in a 1× PBS (Sigma-Aldrich, pH 7.4). The Au nanogrid microelectrode, a platinum (Pt) electrode, and an Ag/AgCl electrode served as the working, counter, and reference electrodes, respectively. EIS measurements were obtained using a 10 mV root mean square (RMS) alternating current voltage with a frequency sweep over a wide range from 100 Hz to 10 kHz. CV measurements were obtained at a scan range of −0.6 to 0.8 V using scan rates from 25 to 300 mV/s. This voltage range falls within the water hydrolysis window. Charge storage capacity (CSC) of the Au nanogrid microelectrodes was calculated at the scan rate of 100 mV/s. For benchtop electrical recording measurements, a Pt electrode input the 10 Hz, 20 mV peak-to-peak amplitude sine wave from a PowerLab 16/35 data acquisition system (ADInstruments Inc.) into a 1× PBS. The Au nanogrid microelectrode was connected to an input channel of the PowerLab data acquisition system to record the sine wave at a 10 kHz sampling frequency. The data was analyzed using a custom MATLAB script.

### Optoelectrical Measurements

A Keithley 2614B source meter (Tektronix Inc.) measured the current-voltage (*I-V*) characteristics of the μ-LEDs and μ-PDs. The responsivities of the optical filter and integrated device were measured under green light illumination from a 527 nm mounted LED light source (M530L3, ThorLabs Inc.). A spectrometer (AvaSpec-ULS2048L, Avantes) monitored the irradiance of the light source and μ-LEDs. The transmittance spectra of the Au nanogrids and optical filters were measured using a spectrophotometer (V-770, Jasco, Inc.). A QE-RS quantum efficiency system (Enlitech) measured the external quantum efficiency (EQE) of the μ-PDs.

### Mechanical Measurements

A motorized test stand (ESM 1500, Mark-10) tested the mechanical flexibility of the probe under different bending cycles.

### Thermal Measurements

A FLIR E8-XT thermal infrared camera monitored the temperature changes of the devices with the μ-LEDs operating under different conditions in air at room temperature.

### Scanning Electron Microscopy (SEM) Measurements

A Raith Pioneer EBL SEM system examined the morphology of the Au nanogrids using a 10 kV acceleration voltage.

## RESULTS AND DISCUSSION

### Optical Characterization of the Microelectrodes

Figure 1c presents a SEM image of the Au nanogrid microelectrodes. The nanogrids have a width of 400 nm and a thickness of 40 nm, respectively. The center to center distance (pitch) between two nearest nanogrids in the network is 6 μm. These nano- and micro-features make it possible to scale down the microelectrode sizes to near cellular-scale (such as the 50 μm diameter in this work) while still maintain an effective conductive network inside the microelectrodes for electrical recording. The nanogrid parameters can be adjusted by EBL (resolution down to sub-10 nm) to fine tune the physical properties of the microelectrodes. The average transmittance values between 400 and 800 nm increase from 71.3 ± 0.42% to 81.5 ± 0.93%, and 88.8 ± 1.1% when the pitch values increase from 4 to 6, and finally 12 μm, respectively (Figure 1d). The average results are over 5 devices unless noted otherwise. The excellent optical transparency in the visible region is beneficial to minimize the crosstalk between electrical recording and optical sensing from both GFRs and fluorescent reporters with red-shifted excitation and emission profiles.

### Electrochemical Characterization of the Microelectrodes

EIS is an important method to study the electrochemical performance at microelectrode/electrolyte interface. Figure 2a presents the EIS results of the 50 μm diameter Au nanogrid microelectrodes with various pitch values. The average electrochemical impedance values at frequencies relevant to biopotential sensing (1 kHz) decrease from 571 ± 38.0 to 319 ± 18.1, and 220 ± 8.44 kΩ with the pitch values decreasing from 12 to 6, and 4 μm, respectively. In all cases, the impedance values are <600 kΩ, indicating that the Au nanogrid microelectrodes are suitable for electrophysiological recording based on the results from Park et al..^24^ The 50 μm diameter Au nanogrid microelectrodes with 400 nm width, 40 nm thickness, and 6 μm pitch are used in the following study unless mentioned otherwise due to its balanced transmittance (81.9% at 550 nm) and impedance (319 kΩ at 1 kHz). Increasing the sizes of the microelectrodes will further reduce the impedance values and allow recording from a larger number of cells at the interface.^30^ The performance of the Au nanogrid microelectrodes is compared to other transparent microelectrodes at similar transmittance levels (>80%), including carbon nanotube (CNT),^36^ graphene,^37^ ITO,^38^ and metal nanostructures^39^ (Figure 2b). The Au nanogrid microelectrodes in the multimodal device exhibit a superior normalized electrochemical impedance (6.3 Ω cm^2^ at 1 kHz), which are among the most competitive performance for flexible transparent microelectrodes currently reported for electrophysiology. The phase spectra in Figure 2c show that the Au nanogrid microelectrodes exhibit more capacitive behaviors (~-75°) at physiologically relevant frequencies from 100 Hz to 1 kHz and gradually become more resistive at higher frequencies (e.g., −55° at 10 kHz).

**Figure 2.**
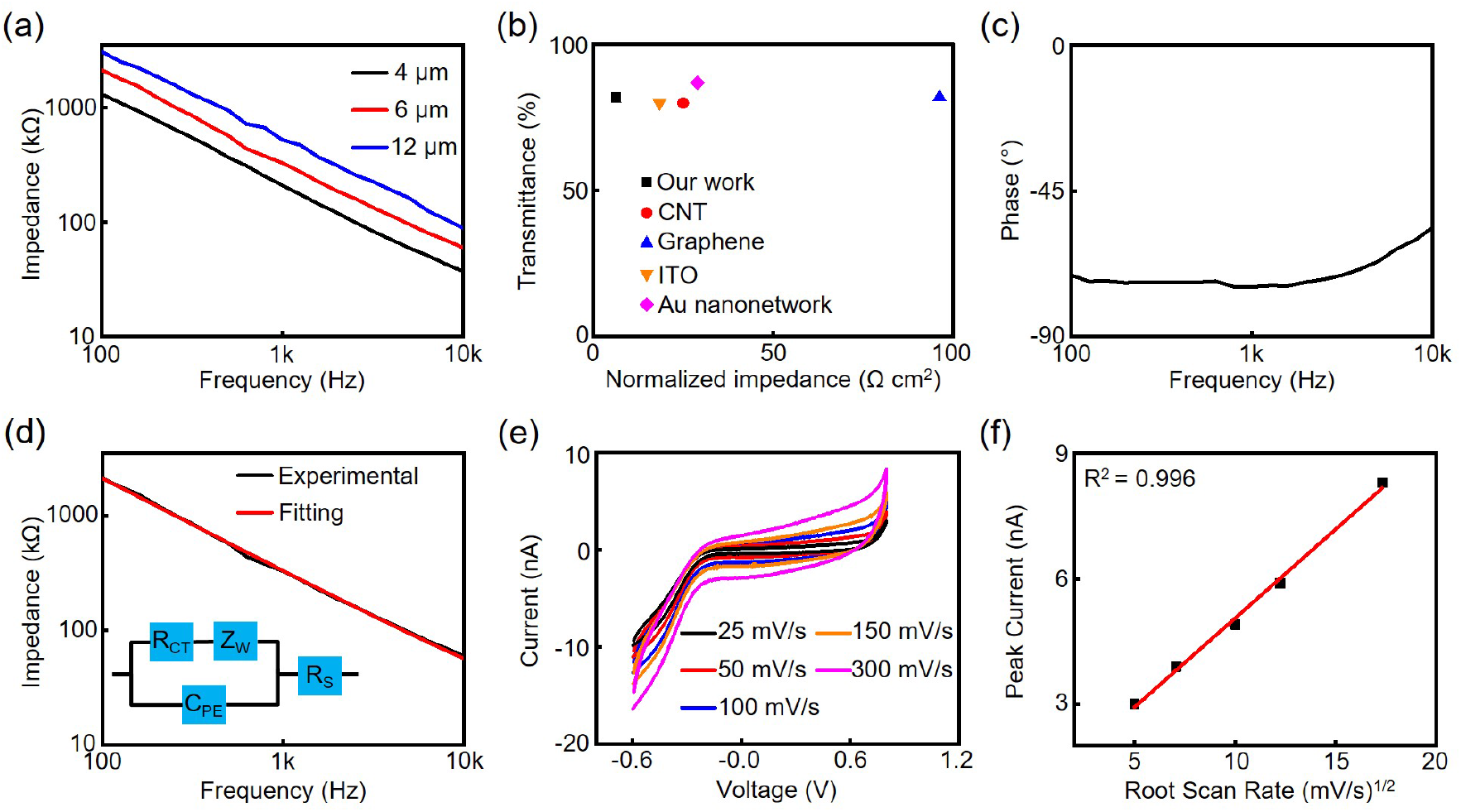
EIS and CV characterizations of the Au nanogrid transparent microelectrodes. (a) Impedance spectra of the 50 μm diameter Au nanogrid microelectrodes with 400 nm nanogrid width, 40 nm thickness, and different pitch values at 4, 6, and 12 μm, respectively. (b) Comparison of optical transmittance and normalized impedance of the Au nanogrid microelectrodes to other reported transparent microelectrodes. (c) Phase spectrum of a 50 μm diameter Au nanogrid microelectrode with 400 nm nanogrid width, 40 nm thickness, and 6 μm pitch. (d) Equivalent circuit model fitting results of the Au nanogrid microelectrodes. (e) CV results of Au nanogrid microelectrodes under scan rates at 25 (black), 50 (red), 100 (blue), 150 (orange), and 300 (magenta) mV/s, respectively. (f) Peak current from CV tests as a function of the root scan rate.

The EIS results are fit to an equivalent circuit model to provide more insights into the electrochemical characteristics of the Au nanogrid microelectrodes (Figure 2d).^19^, ^30^ The model includes a constant phase element (*C_PE_*) in parallel with a charge transfer resistance (*R_CT_*) and a Warburg impedance for diffusion (*Z_W_*), and a solution resistance (*R_S_*). The *C_PE_* is defined by 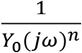. Here *Y_0_* is the *C_PE_* magnitude, *ω* is the angular frequency, *j* is the unit imaginary number, and *n* is a constant from 0 to 1. The large *n* value (0.96) from the modeling indicates that the Au nanogrid microelectrodes in the multimodal device exhibit a more capacitive interface similar to an ideal capacitor (*n* = 1), which is desired for electrical recording applications.

CV measurements examine the microelectrode/electrolyte interfaces and potential electrochemical reactions on the microelectrodes. Figure 2e presents CV data of the Au nanogrid microelectrodes at different scan rates from 25 to 300 mV/s, respectively. No oxidation/reduction peaks are observed. The linear dependence of the peak currents on the square root of scan rate from the CV results (Figure 2f) indicates a diffusion-controlled electron transfer process at the nanogrid/electrolyte interface.^40–41^ The microelectrodes exhibit an average CSC of 2.28 ± 0.2 mC/cm^2^, comparable to alternative transparent microelectrodes.^42–43^

### Optoelectrical Characterization of the Optical Sensing Subsystems

To measure fluorescence with on-chip μ-PDs, it is crucial to directly integrate effective optical filters on the μ-PDs to prevent the μ-LED fluorescence excitation light from reaching to the nearby μ-PDs. This is because the excitation light typically has an irradiance that is several orders of magnitude higher than that of the resulting fluorescence signals. We have previously designed optical filters with absorber dyes mixed in polymers to block light from blue μ-LEDs.^21–22^ Here we optimize the performance of the optical filter by adjusting the concentrations (2.0, 2.5, and 3.0 wt%) of the absorber dye dissolved in a 7 μm transparent SU-8 layer (Figure 3a). The emission peaks of GFRs are normally between 510 and 520 nm. As expected, the transmittance ratio of the light at 520 to 462 nm (maximum μ-LED emission wavelength) increases from 17.2 to 99.1, and finally 655 when the concentration of the absorber dye increases from 2.0 to 2.5, and 3.0 wt%, respectively. The color of the optical filters darkens from light green to dark green with increased dye concentration (Figure 3b), which is in line with the transmittance results. Although higher rejection of the excitation light is desired to minimize the crosstalk, the optical filter with 2.5 wt% of the absorber dye is used as the balanced optical filter condition due to its moderate transmittance in the green region (~40% at 520 nm) so that as much fluorescence emission light as possible could still pass the optical filter and be captured by the μ-PD. Importantly, integrating the Au nanogrids on the optical filter results in negligible changes in the transmittance ratio (Figure 3c) due to the high and uniform transmittance of the Au nanogrids in the visible region.

**Figure 3.**
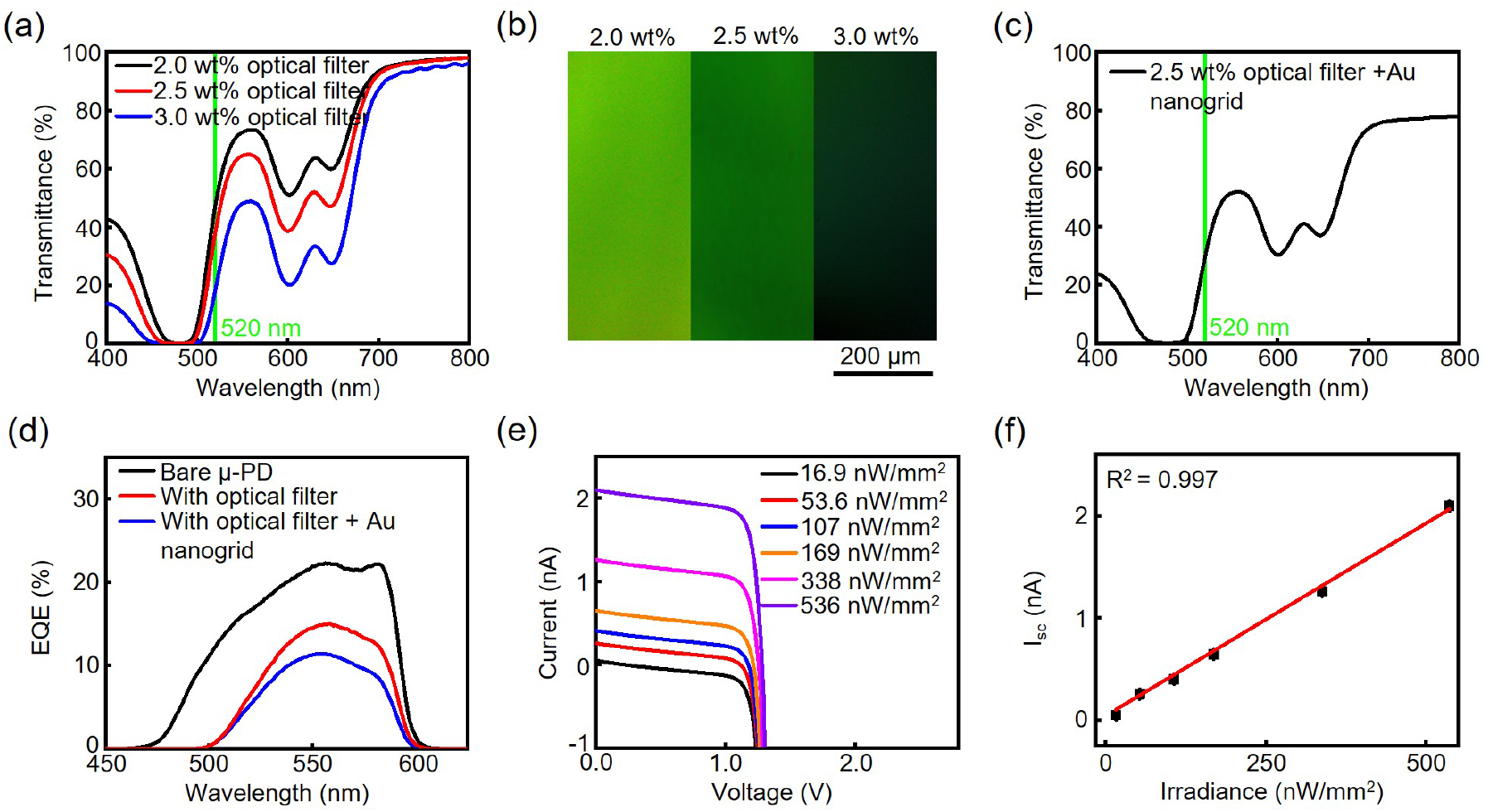
Optoelectrical characteristics of the opto-electric devices. (a) Transmittance spectra of the SU-8 2007 optical filter with 2.0 (black), 2.5 (red), and 3.0 (blue) wt% absorber dye concentration. (b) Optical images of the optical filters at different absorber dye concentrations. (c) Transmittance spectrum of the SU-8 2007 optical filter with 2.5 wt% absorber dye concentration and the Au nanogrid microelectrode ontop. (d) EQE spectra of a bare μ-PD (black), a μ-PD coated with the optical filter (red), and a μ-PD integrated with both the optical filter and the Au nanogrid microelectrode ontop (blue), respectively. (e) Light irradiance-dependent *I-V* characteristics of the μ-PDs under green LED illumination. Black, red, blue, orange, magenta, and violet represent the light irradiance of 16.9, 53.6, 107, 169, 338, and 536 nW/mm^2^, respectively. (f) Light irradiance-dependent *I_sc_* of the μ-PDs under different green irradiance in (e).

The optoelectrical performance of the μ-PDs in the multimodal devices is first characterized using EQE measurement (Figure 3d). The bare μ-PDs show higher EQE values in the green region than the blue region. The EQE values after integrating the optical filter and nanogrids are 4.96 × 10^-5^ and 6.53 × 10^-2^ at 462 and 520 nm, respectively. The EQE ratio at 520 to 462 nm (this ratio defines the wavelength selectivity towards recording green fluorescence from the device) is 1.32 × 10^3^, which is 20 and 12 times higher than those of our previously reported single function optical recording systems.^21–22^ The high wavelength selectivity will minimize the influence of excitation light on fluorescence signals. Figure 3e presents the *I-V* curves of the integrated μ-PDs under 527 nm green light from 16.9 to 536 nW/mm^2^, which cover the typical fluorescence irradiances of GFRs. The short-circuit current (*I_sc_*) is plotted as a function of irradiance, shown in Figure 3f. The integrated μ-PDs have a near-precise linear response (R^2^ = 0.997) to irradiances in this range, suggesting their capability to measure GFR emissions.

### Long-term Stability and Mechanical Flexibility of the Opto-electric Device

Stable device operation when immersed in biofluids is imperative for biointegration. A soaking test in a 1× PBS (pH 7.4) at 37 °C simulates the body fluid environment and evaluates the durability and stability of the opto-electric devices for long-term implantable applications. All components are soaked without operation. The *I_sc_* of the μ-PDs in the dark, 1 kHz electrochemical impedance values from the Au nanogrid microelectrodes (Figure 4a), and *I-V* characteristics of the blue μ-LEDs (Figure 4b) show negligible variations before and after soaking in PBS for up to 40 days. The results indicate the opto-electric devices are reliable for chronic use. Mechanical flexibility is another important feature for implantable devices to minimize tissue damage, inflammation, stress at device/tissue interface and minimize device performance degradation overtime after implantation.^44–45^ Figure 4c,d shows that the *I_sc_* of the μ-PDs in the dark, the irradiances from the μ-LEDs, and the 1 kHz electrochemical impedance values from the Au nanogrid microelectrodes in the opto-electric devices remain stable after 5,000 bending cycles against a small radius of 5 mm. This radius is comparable to the anatomical features of interest in small animals.^37^ Together, those results indicate the excellent chronic stability and mechanical compliance of the opto-electric devices.

**Figure 4.**
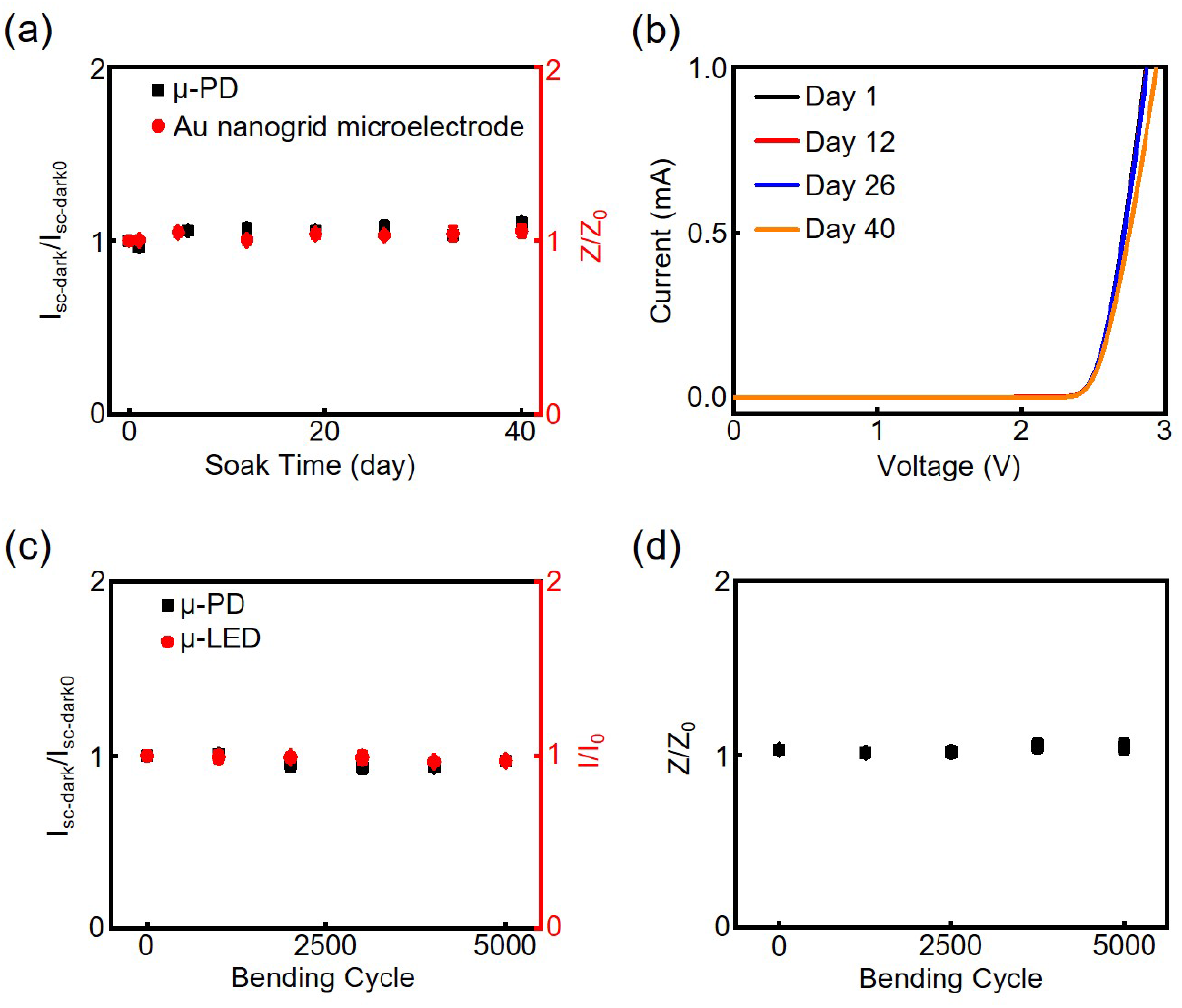
Long-term stability and mechanical flexibility of the opto-electric devices. Changes of *I_sc_* of the μ-PDs in the dark and 1 kHz impedance from the Au nanogrid microelectrodes (a), and *I-V*characteristics from the μ-LEDs (b) as a function of soaking time in a 1× PBS at 37 °C. *I_sc_* of the μ-PDs in the dark and μ-LED irradiances (c), and 1 kHz Au nanogrid microelectrode impedance (d) as a function of bending cycle at a radius of 5 mm. I_sc-dark_, Z, and I are the I_sc_ in the dark, impedance, and irradiance at a specific bending cycle. I_sc-dark0_, Z_0_, and I_0_ are the initial I_sc_ in the dark, impedance, and irradiance, respectively.

### Thermal Property of the Opto-electric Device

Tissue damage caused by the heat produced from the μ-LEDs during optical recording is a possible concern for future *in vivo* applications. The maximum temperature changes of the opto-electric devices are investigated in air with the blue μ-LEDs operating at irradiances from 10 to 20 mW/mm^2^, duty cycles from 0 to 90%, and frequencies from 0.5 to 50 Hz. Those conditions are relevant to optical fluorescence recording.^21, 46^ As expected, temperature increases are larger at higher irradiances (Figure 5a). Increasing the duty cycles results in larger temperature changes due to the reduced time to dissipate the accumulated heat on the device surface at higher duty cycles. Figure 5b shows that changes in temperature are smaller at higher frequencies. Overall, the maximum temperature increases from the opto-electric devices are well below the thermal thresholds (2 °C)^47^ for tissue damage when the blue μ-LEDs in the opto-electric devices operate at the conditions in Figure 5a,b. Importantly, the temperature increases *in vivo* are likely to be much lower than those measured in air due to the faster heat dissipation in biofluids.

**Figure 5.**
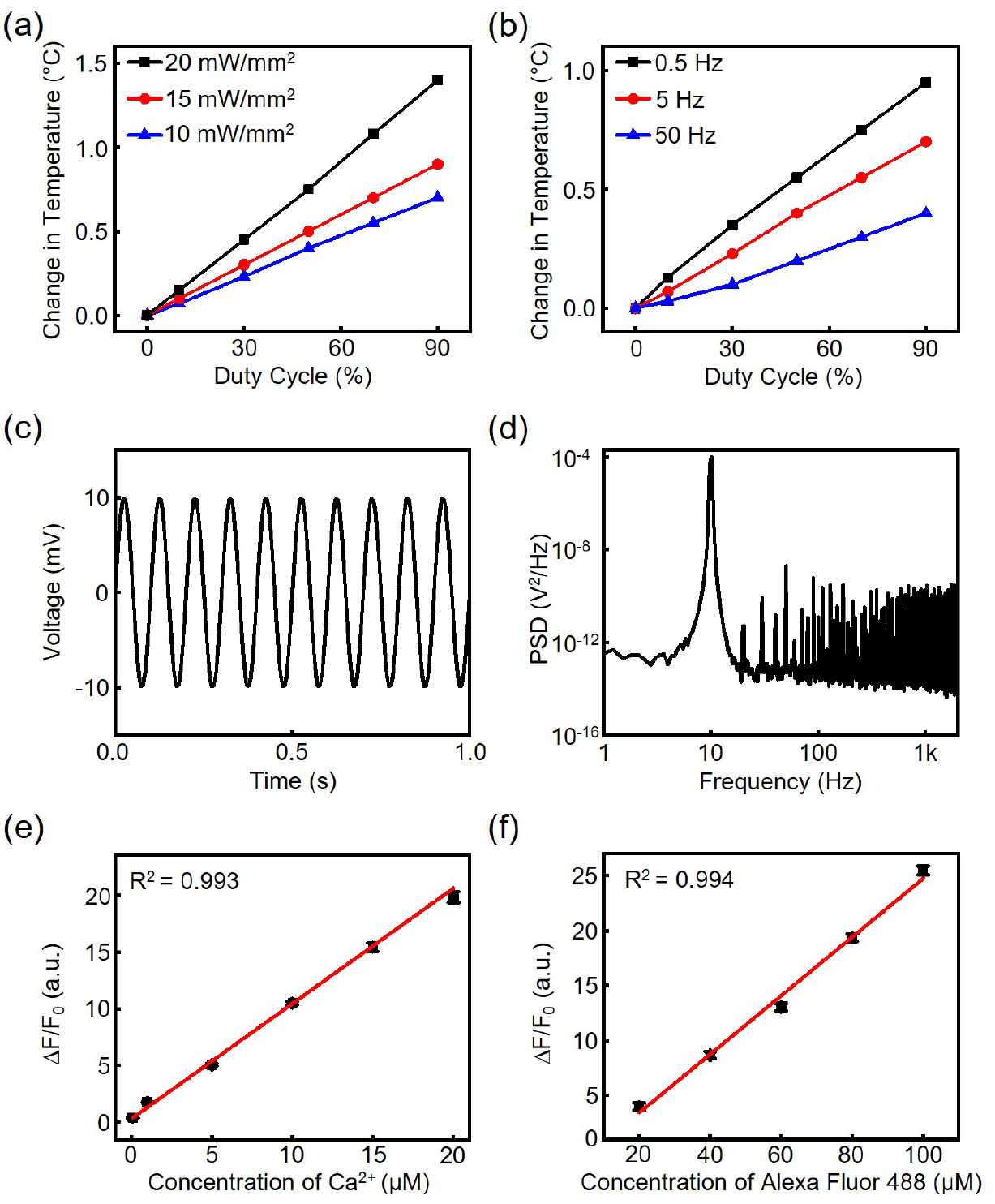
Benchtop measurements of the opto-electric devices. Temperature change versus duty cycle from the μ-LEDs in the opto-electric devices in air at (a) 5 Hz with 10, 15, and 20 mW/mm^2^ output irradiance and (b) 0.5, 5, and 50 Hz with 10 mW/mm^2^ output irradiance, respectively. (c) Representative electrical recording output of a programmed 10 Hz, 20 mV peak-to-peak sine wave input by the Au nanogrid microelectrode in the flexible multimodal opto-electric device in a 1× PBS. (d) Representative PSD graph of the Au nanogrid microelectrode recorded sine wave output in (c). Fluorescence emission intensity results from Oregon Green 488 BAPTA-1 calcium dye solutions with different calcium concentrations from 0.1 to 20 μM (e), and Alexa Fluor 488 dye solutions with concentrations from 20 to 100 μM (f), respectively.

### Benchtop Recording of Electrical Waves and GFRs

The recording capability of the Au nanogrid microelectrodes in the opto-electric devices is tested in a 1× PBS. Figure 5c presents the recorded signals of a programmed 10 Hz, 20 mV peak-to-peak amplitude sine wave input. No observable decrease in signal amplitude occurs. The power spectral density (PSD) provides insights into the noise and signal in the frequency domain (Figure 5d). The PSD results feature a peak at 10 Hz from the input signals. The signal-to-noise ratio and RMS noise are 40.3 dB and 62.5 μV, respectively.

Intracellular calcium levels span from ~0.1 μM when cells (e.g., neurons and cardiomyocytes) are at rest to 10-20 μM under stimulation.^9^, ^48^ To demonstrate the capability of the devices for recording intracellular calcium dynamics, fluorescence emission intensity in this concentration range is measured using Oregon Green 488 BAPTA-1 (excitation/emission maxima at 494/523 nm) as the calcium indicator (Figure 5e). A superior linearity in response (R^2^ = 0.993) is observed in the fluorescence signals. In addition to calcium measurements, the devices can record fluorescence from other GFRs. For example, Figure 5f presents the fluorescence recording results from a widely used GFR in imaging/labeling, Alexa Fluor 488 dye (excitation/emission maxima at 490/525 nm). The concentrations of Alexa Fluor 488 dye are from 20 to 100 μM, which are consistent with those used *in vivo*.^49–51^ The fluorescence signals increase linearly with increased Alexa Fluor 488 dye concentrations (R^2^ = 0.994). Together, those results show the various fluorescence intensity changes associated with cell functions from GFRs are within the dynamic range of the opto-electric devices.

## CONCLUSIONS

The flexible and multimodal opto-electric devices reported here address the current technical limitation of integrating transparent microelectrodes ontop of microscale light sources and photodetectors for precise co-localized optical fluorescence and electrical recording of cell activity. The unique monolithic fabrication process is advantageous to combine various microscale optical and electrical components on thin flexible substrates for advanced implantable applications. The ultrathin nature of the transparent microelectrodes results in a negligible increase in the dimensions of the multimodal devices compared to existing single function optical recording devices. The microelectrodes exhibit a small diameter of 50 μm, a high transmittance >81% in the visible region, and a low normalized impedance of 6.3 Ω cm^2^ for high-fidelity recording of electrophysiological signals. The optical recording subsystem features a high wavelength selectivity >1,300 and a high linearity (R^2^ >0.99) for measuring green fluorescence from GFRs at conditions mimicking those *in vivo*. The resulting multimodal systems exhibit superior chronic stability during a soak test in a 1× PBS for 40 days, and durable mechanical flexibility under cyclic stretching of 5,000 cycles. In addition, thermal properties of the devices are carefully characterized to identify operating parameters that avoid thermal damage to the surrounding tissue. Further directions include (1) increasing the number of recording channels in each modality to spatially probe multiple cell/tissue/organ regions; (2) expanding the optical recording capability to red-shifted fluorescent reporters; (3) developing miniaturized systems for fully wireless operations; (4) integrating actuator functionality to realize closed-loop physiological investigations. Collectively, these advancements will open up new opportunities in chronic multimodal recording *in vivo*, and provide essential diagnostic and therapeutic information for different forms of organ dysfunction and disease, which are beyond the capability of current optical fluorescence and electrical recording tools.

## Notes

The authors declare no competing financial interest.

## ACKNOWLEDGMENT

This work was finacially supported by National Science Foundation, grants number 2011093 and 2131682; and The George Washington University Cross-Disciplinary Research Fund. The authors thank The George Washington University Nanofabrication and Imaging Center for its facilities and support for device fabrication.

